# Absorbable Conductive Electrotherapeutic Scaffolds (ACES) for Enhanced Peripheral Nerve Regeneration and Stimulation

**DOI:** 10.1101/2022.07.30.500547

**Authors:** Shriya Srinivasan, Lisa Gfrerer, Paramesh Karandikar, Avik Som, Amro Alshareef, Sabrina Liu, Haley Higginbotham, Keiko Ishida, Alison Hayward, Sanjeeva P. Kalva, Robert Langer, Giovanni Traverso

**Author notes:** Please address correspondence to, (GT) and (SS).

## Abstract

While peripheral nerve stimulation (PNS) has shown promise in applications ranging from peripheral nerve regeneration after injury to therapeutic organ stimulation, clinical implementation has been impeded by various technological limitations, including surgical placement, lead migration, and atraumatic removal. Here, we describe the design and validation of a new platform for nerve regeneration and interfacing: Absorbable, Conductive, Electrotherapeutic Scaffolds (ACES). ACES are comprised of an alginate/poly-acrylamide interpenetrating network hydrogel optimized for both open and minimally invasive percutaneous approaches. In a rodent model of sciatic nerve repair, ACES significantly improved motor and sensory recovery (*p* < 0.05), increased muscle mass (*p* < 0.05), and increased axonogenesis (*p* < 0.05). Triggered dissolution of ACES enabled atraumatic, percutaneous removal of leads at forces significantly lower than controls (*p* < 0.05). In a porcine model, ultrasound-guided percutaneous placement of leads with an injectable ACES near the femoral and cervical vagus nerves facilitated stimulus conduction at significantly greater lengths than saline controls (*p* < 0.05). Overall, ACES facilitated lead placement, stabilization, stimulation and atraumatic removal enabling therapeutic PNS as demonstrated in small and large animal models.

## Introduction

Peripheral nerve injury or dysfunction critically affects somatic and autonomic function, resulting in pain, immobility, and/or loss of functionality, which can significantly reduce quality of life(*1, 2*). With over twenty million annual cases worldwide, peripheral nerve injuries result in an annual economic burden of $150 billion USD(*3, 4*). However, less than half of cases yield satisfactory motor or sensory recovery after receiving standard medical care primarily due to poor axonal regeneration and disuse atrophy of distal muscles(*1*),(*5*). Moreover, injuries and neuropathies occurring in autonomic nerves or in anatomically challenging locations, such as the celiac plexus, remain difficult to cure and are treated largely through non-specific pharmacotherapies(*6*).

### The promise of PNS

Electrical peripheral nerve stimulation (PNS) has been clinically demonstrated to significantly improve sensory and motor function in contexts where surgery alone has had limited effectiveness(*7*–*9*)^,1^. Following nerve crush, transection or stretch injury, PNS accelerates axonogenesis by modulating plasticity, elevating neuronal cyclic adenosine monophosphate, and upregulating neurotrophic factors and Schwann cell activity(*9*). Chronic PNS prevents disuse atrophy of distally innervated muscles by reducing apoptosis of denervated muscle fibers, which preserves muscle mass. Moreover, for autonomic conditions of neural etiology, such as incontinence(*10*), sleep apnea(*11*), and gastrointestinal motility(*12*), PNS can restore and even augment function (Figure 1a). Despite the widespread and evolving evidence of the benefits of PNS, its clinical implementation has been limited to treating intractable pain pathologies such as lower back pain and occipital neuralgia(*13*); little to no clinical translation of PNS has occurred for peripheral nerve repair(*1, 7, 14, 15*) or for non-pain indications in autonomic nerves (Figure 1a)^1,2^ due to hardware implantation challenges.

**Figure. 1.**
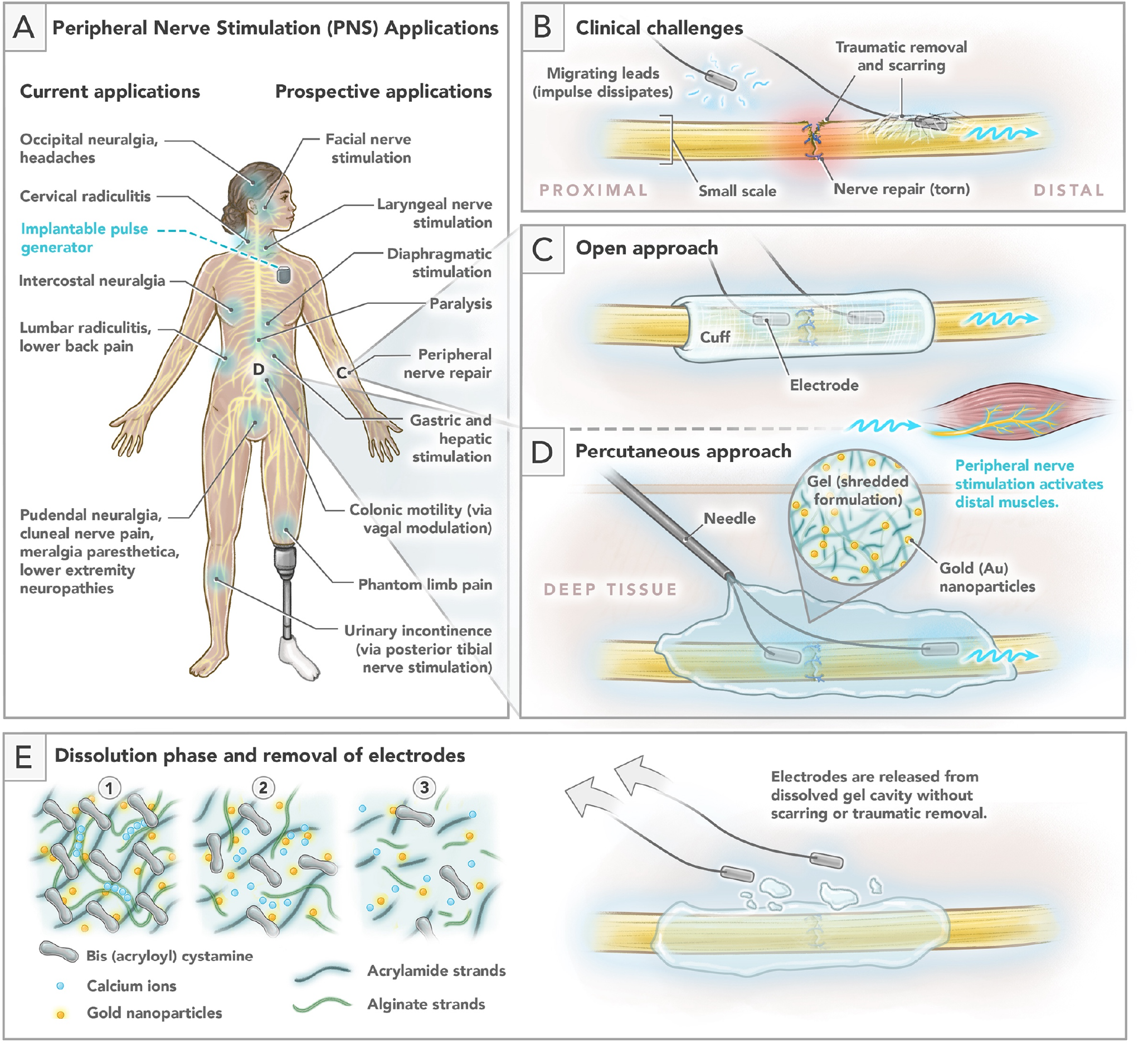
Absorbable Conductive Electrotherapeutic Nerve Scaffold (ACES) for Peripheral Nerve Stimulation. A) Current and prospective applications for peripheral nerve stimulation. B) Clinical challenges of implanted peripheral nerve stimulation hardware include migration, scarring, precise targeting of millimeter caliber nerves and traumatic removal. C) In an open approach, a cuff embodiment of ACES stabilizes electrodes on the proximal and distal sides of a repaired nerve repair. Stimulation propagates to the distal muscle. D) In a percutaneous approach, the injectable embodiment of ACES stabilizes electrodes and conducts stimulation signals from the leads to the nerve. E) ACES dissolution occurs through chelation of calcium, releasing cross-links. Once the scaffold is cavitated, leads can be atraumatically removed with minimal force.

### The limitations of PNS hardware implantation

Leads for peripheral nerves are commonly placed using an open surgical approach (e.g., to treat extremity nerve injuries, or for migraine surgery), while autonomic nerves deep in the body (e.g., in the celiac plexus, spinal cord, or SMA plexus adjacent to the aorta) are accessed through minimally invasive percutaneous approaches including laparoscopy, robotics, and interventional radiology. Placement of nerve stimulation hardware is complicated by the need to precisely target millimeter-sized nerves to prevent off-target stimulation^4^, achieve lead stabilization(*16*) on delicate neural tissue, and avoid mechanical interference with nerve repairs (Figure 1b). Following implantation, lead migration is a common complication, with 60-100% of cases requiring subsequent re-operation^4,5^ (as per the Manufacturer and User Facility Device Experience (MAUDE) database, Supplemental Note 1). Atraumatic removal is complicated by tissue adhesion to the leads. Consequently, at the completion of therapy or when the device malfunctions, leads are frequently left in the body, posing potential discomfort, and serving as a nidus for infection. While bioadhesive systems have been previously developed, the lack of triggerable dissolution and biocompatibility limit their utility as an electrode scaffold. Some systems require in vivo free radical polymerization which can present a potential carcinogenic risk(*17*–*19*). As a result, PNS clinical trials for nerve repair have been constrained to intra-operative or one-hour post-operative(*20*),(*21*) periods. A conformable scaffold capable of stabilizing leads, bridging conduction gaps between electrodes and target nerves, and releasing hardware on demand for atraumatic removal is necessary for the clinical implementation of PNS.

### Absorbable Conductive Electrotherapeutic Scaffolds (ACES)

Here, we describe the design and validation of an **absorbable, conductive, electrotherapeutic scaffold (ACES)** *(Figure 1)*. Comprised of an alginate/poly-acrylamide-based interpenetrating network (IPN) hydrogel impregnated with gold nanoparticles, the scaffold facilitates placement and stabilization of leads at target sites by mechanically conforming to the surrounding anatomical features. ACES restricts granulation and scar formation to the exterior of a dissolvable gel cavity, rather than around the electrodes, to maximize stimulation efficacy and minimize trauma upon removal. When therapeutic stimulation is complete and the hardware is to be removed, the scaffold can be triggered to dissolve, facilitating atraumatic removal. ACES can be designed in application-specific embodiments. For nerve repairs utilizing an open surgical approach, ACES can be preformed into a cuff, situating one electrode each on the proximal and distal sides of a nerve repair site to provide electrical stimulation (ES) for axonogenesis and distal organ stimulation *(Figure 1c)*. Alternatively, a grounded-gel formulation of ACES can be applied for percutaneous and open surgical approaches. This formulation can be injected at the site of the nerve, where it stabilizes electrodes, conforms to the surrounding anatomical features, and establishes a conductive path from the leads to the target nerve without requiring direct electrode-nerve contact *(Figure 1d)*. Upon triggered dissolution through a co-implanted microcatheter or injection, electrodes are released from the gel into the cavity of the scaffold and can be removed transcutaneously in an atraumatic fashion (Figure 1e). The gel’s mechanical properties are matched to that of the nerve to permit nonintrusive support, prevent rejection, and provide mechanical support(*22*), accelerating axonogenesis.

In this study, we test the features of the ACES in an open surgical model of sciatic nerve transection and in a minimally invasive percutaneous femoral/vagal nerve stimulation model to evaluate its efficacy. In comparison to untreated controls, we hypothesize that ACES will facilitate lead placement, mitigate migration and improve axonogenesis, motor and sensory function. We further hypothesize that the triggered dissolution of ACES will enable the removal of implanted electrodes with less force than intact scaffolds, minimizing tissue trauma. We then evaluate the conductive ability of the gel to bridge the anatomical distances between implanted electrodes and the target nerves in which stimulation is possible.

## Results

### Design rationale

To enable optimal tissue adherence and trauma-free removal of leads, we designed a tough, adhesive polymer that could undergo a significant decrease in strength after dissolution. Covalently crosslinked polyacrylamide was selected for its adhesive and elastic properties while ionically crosslinked alginate was selected for its flexibility and triggerable weakening(*23*). To enable triggerable dissolution of the polyacrylamide network, N,N’-Bis(acryloyl)cystamine was chosen as the covalent crosslinker due to its labile disulfide bond (Supplemental Figure 8). To optimize the IPN formulation, various combinations of acrylamide, alginate, and crosslinker concentrations were evaluated for adhesive strength and subsequent dissolution capacity through an electrode removal assay (Methods). The elastic modulus of the gel was tuned to be comparable to peripheral nerve (772.8 ± 244.3 - 4387.6 Pa(*24*)). A ratio of 50mg/mL acrylamide and 25 mg/mL alginate with 75% crosslinker concentration resulted in the greatest reduction in strength after dissolution (Figure 2a,b). Upon adhesion of the IPN to the electrodes, the leads withstood up to 4.96 +/- 0.08 N of tensile force with no measurable displacement. Dissolution was initially performed with 0.5 M reduced glutathione (GSH), which cleaves the labile disulfide bond within the bis(acryloyl)cystamine crosslinker, and EDTA (Ethylenediaminetetraacetic acid), which chelates Ca^2+^ ions and thus cleaves the ionic alginate crosslinks. Upon dissolution, the electrodes slid out with 2.3 +/- 0.03 N of force. To evaluate the time-dependent viscoelastic behavior, an oscillatory shear test was performed before and after the dissolution of ACES (Figure 2c). As anticipated, the storage and loss moduli significantly decreased following dissolution (> 3 fold, *p* < 0.01, Student’s t-test). The reduction in loss modulus reflects the dissolved alginate crosslinks, which allow the leads to slide out with little frictional force. To determine the optimal incubation time for dissolution, we rheometrically characterized the storage and loss moduli of the polymer under immersion for a period of 24 hours. At 30 minutes, an 85% reduction in storage modulus and 98% reduction in loss modulus was observed (Figure 1d). Within 1 hour, polymer weakening was comparable to that of 24 hours. Thus, 60 minutes was determined to be sufficient for dissolution. To further optimize the dissolution profile, we tested a variety of formulations of GSH, SBC (sodium bicarbonate), and EDTA at 30- and 60-minute periods using an electrode removal assay. All dissolution treatments enabled electrode removal at the pre-strain threshold of the tensile test (0.01N), requiring no additional force to pull out the electrode (Figure 2e). In contrast, in controls with no treatment, 7.62N of force was required to pull out the electrode—an over 750-fold increase in force. A solution of 0.5M GSH+ 0.5M SBC + 1M EDTA was selected for its ability to effectively dissolve the ACES.

**Figure 2.**
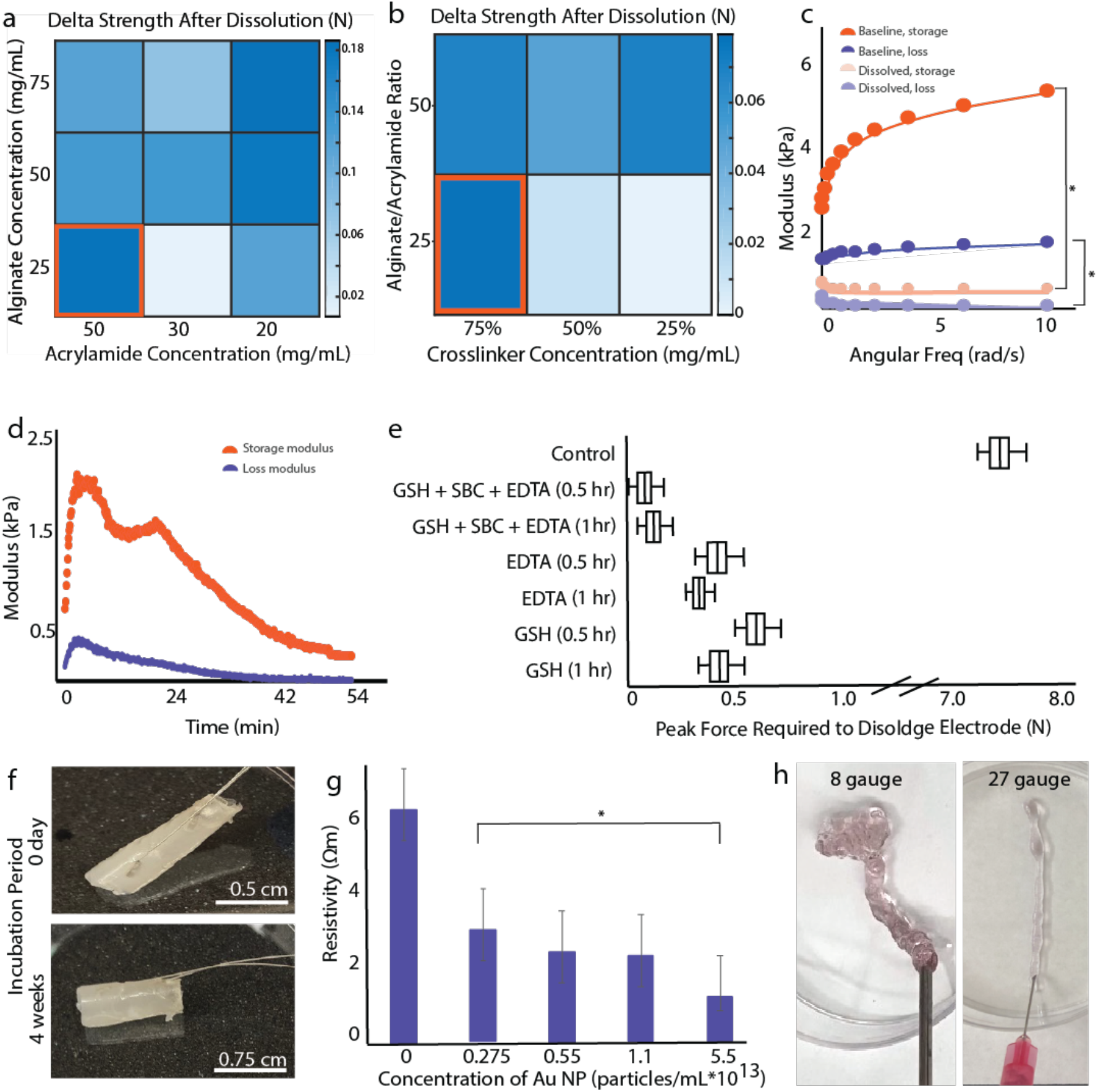
Optimization of triggerable polymeric formulation for ACES. a) Optimization of acrylamide and alginate concentrations yielded the greatest differential strength in the 50mg/mL acrylamide and 25 mg/mL alginate formulation (orange). b) Optimization of the crosslinker concentrations yielded the greatest differential strength with 75 ug/mL for a 25% c) Storage and loss moduli before and after dissolution demonstrate significant weaking of the ACES following dissolution. d) Time course of dissolution demonstrates a monotonal decrease in the storage and loss moduli of the scaffold polymer. e) Comparison of various dissolution formulations. f) Appearance of the ACES in a cuff embodiment at 0 and 4 weeks of incubation in saline. g) Resistivity of ACES formulations. h) The conformable embodiment of ACES can be extruded through a range of needle gauges from 8 – 34. For all panels, measurements represent the average of three independent samples.

For open surgical approaches, a preformed cuff-like scaffold shape provides mechanical support and eases surgical placement around the nerve(*25*). Thus, ACES was preformed into a custom mold, incorporating electrodes and a microfluidic channel to deliver the dissolution reagents (Figure 2f top). Following a four-week incubation in saline at 37°C to mimic the tissue environment, no significant degradation, tearing or disintegration was observed (Figure 2f bottom). Furthermore, the ACES’s modulus and electrode removal assay results before and after incubation demonstrated no significant differences (*p* < 0.05, Student’s two-tailed heteroscedastic t-test, n = 4 trials).

To bridge the conductivity gap between stimulation hardware and the target nerve in minimally invasive surgical approaches, the conductivity of the scaffolding material was increased by incorporating gold nanoparticles (Au NPs, 40nm), selected for their biocompatibility and conductivity. Compared to ACES with no added Au NPs, sheet resistance decreased 5-fold with an Au NP concentration of 5.5×10^13^ particles/mL as per a four-point contact test (Methods, Figure 2g). The gel’s self-healing properties(*26*) were leveraged to create an injectable embodiment. After polymerizing ACES as a 2mm sheet, thin 2-4mm strips were cut and loaded into a syringe. Injectability was then determined using extrusion equations(*27*). Given the gel’s consistency index of K = 21.996 Pa*s^n^ and a shear-thinning index (n) of 0.134, it was determined that the gel could be injected through needles ranging from 7 – 34 gauge under a pressure of 2.6 MPa (human force of 50 N on a 5mm diameter plunger) using the power-law fluid equations; these calculations were also validated empirically (Figure 2h, Supplemental Figure 1).

The biocompatibility of the ACES system was evaluated using an extract exposure test(*28*) at treatment concentrations between 50 – 200 mg/mL and with varying Au NP concentrations at 1 and 7 days. Cell viability normalized to the vehicle treatment group was greater than 100% after 24 hours and greater than 70% after 7 days, which is considered non-toxic by ISO 10993 norms(*28*) (Supplemental Figure 2).

### ACES in open surgical repair of sciatic nerve transection

Lewis rats (450-500g, 10-12 weeks old) were randomly assigned to 1) a control group (n= 9), receiving the ACES cuff without PNS or dissolved removal, 2) an experimental group (n=9) receiving the ACES cuff, PNS three times per week, and dissolved removal or 3) an experimental group receiving injectable ACES without PNS.

In all groups, the sciatic nerve was transected, acutely repaired, and supported by the ACES (Figure 3a-e) in either cuff or injectable embodiments, requiring less than 5 or 2 minutes for stable placement, respectively (Supplemental Movie 1,2). Over the course of 6 weeks, electrical stimulation applied at the sciatic nerve resulted in selective activation of the hind limb; no significant change in electrode impedance was observed, suggesting no lead migration in animals with ACES. Six weeks postoperatively, compound muscle action potentials (CMAPs) in the experimental group were significantly higher than untreated controls (*p* < 0.05, Student’s two-tailed heteroscedastic t-test) with a significantly lower stimulation-response threshold (*p* < 0.05, Student’s two-tailed heteroscedastic t-test) (Figure 3f). The sensory thresholds in the control animals were significantly higher in the affected limbs compared to the contralateral unaltered limbs (*p* < 0.05, Student’s two-tailed heteroscedastic t-test), suggesting incomplete sensory reinnervation (Figure 3g). In contrast, the sensory threshold ratio between the affected limbs and unaltered limbs in the experimental group was 1.02, suggesting a full sensation recovery in the limb supported by the ACES-based rehabilitation. The sensory thresholds of the contralateral unaffected sides of animals in each group were insignificantly different (*p* < 0.001, Student’s two-tailed heteroscedastic t-test). Following euthanasia, the ratio of mass of the explanted gastrocnemius (GSC) and tibialis anterior (TA) muscles were compared to their contralateral controls to assess the impact of ACES on minimizing disuse atrophy. Animals with stimulation facilitated by ACES demonstrated significant reductions in disuse atrophy as compared to their controls (*p* < 0.05, Student’s two-tailed heteroscedastic t-test, Figure 3h,i). To quantify axonal regeneration, axon counts were performed in the proximal and distal segments of the sural, tibial, and peroneal nerves harvested distal to the site of repair (Figure 3k). On the contralateral side, the proximal-distal axon count ratios were approximately one and not significantly different between control and experimental animals. However, on the affected side, the proximal-distal axon count was significantly higher in the experimental group in the sural and peroneal nerve (*p* < 0.05, Student’s two-tailed heteroscedastic t-test, Figure 3k). These results suggest that ACES facilitates lead stabilization, improves axonogenesis, and supports chronic PNS which, in turn, yields functional recovery and prevents disuse atrophy.

**Figure 3.**
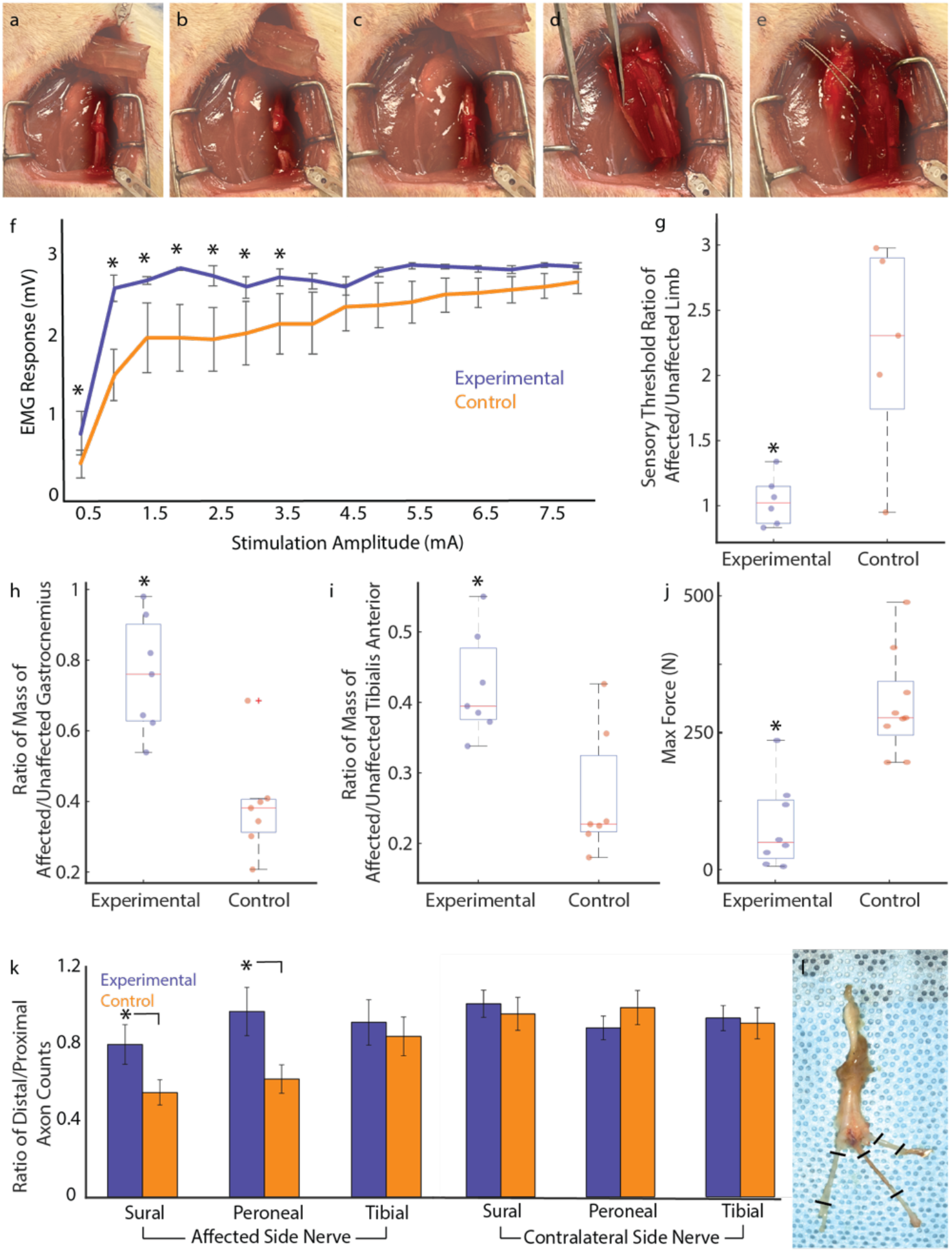
ACES in open surgical repair of sciatic nerve transection. Surgical process involved the a) exposure of the sciatic nerve, b) transection of the sciatic nerve proximal to the trifurcation c) microsurgical coaptation of the sciatic nerve, d) surgical placement of ACES around repair site e) and stabilization and visualization of the leads in the scaffold. f) EMG response of the tibialis anterior was measured in response to proximal sciatic nerve stimulation via the ACES electrodes resulting in a significantly higher EMG response in the experimental group than the control at stimulation amplitudes between 1 – 3.5mA. g) The ratio of the sensory threshold in the affected to unaffected limb in the control group was significantly higher than the experimental, stimulated group (p < 0.05). h) The ratio of mass of the affected to unaffected gastrocnemius was significantly higher in the experimental group than the control group (p < 0.05). i) The ratio of mass of the affected to unaffected tibialis anterior was significantly higher in the experimental group than the control group (p < 0.05, n = 9/group). j) The maximum force (N) needed to remove the electrodes was greater in the control group than the experimental group. k) The ratio of the distal to proximal axon counts were calculated in the sural, peroneal, and tibial branches on the affected side and compared to the contralateral side. The experimental group had a significantly higher distal-proximal axon count ratio in the sural and peroneal branches on the affected side. l) Image of the sciatic nerve descending into the three branches of sural, peroneal, and tibial. Hashes indicate the locations of axon counts.

To characterize ACES-based hardware removal, the force required to extract the implanted electrodes was measured with and without triggered dissolution. In dissolved ACES cuffs, the peak force and tensile stress in the surrounding tissue were significantly lower (*p* = 0.005 and *p* = 0.0069 respectively, n = 9/group) as compared to non-dissolved controls (Figure 3j, Supplemental video 1). In animals with dissolved injectable ACES, the force required for removal was significantly lower (*p* < 0.04, two-tailed heteroscedastic t-test) at 1.66 +/- 0.9N while undissolved controls required 5.87 +/- 2.9N. In cases with undissolved scaffolds, significant stretching and tearing through tissue layers was observed as the electrodes were extracted. Dissection to the implant site revealed little to no trauma of the repaired nerve with dissolved ACES, while tearing of adhesions to the nerve and in two cases, a disrupted nerve repair was observed in the control (Supplemental Figure 3). Upon explant, adhesions to the explanted electrodes were quantified by a blinded plastic surgeon (0 – no adhesion, 5 – fully adhered to scar tissue). Those induced with the dissolution of ACES scored 0.2 +/- 0.44 while the controls scored 4.8 +/- 0.44 for both cuff and injectable embodiments of ACES (Supplemental Figure 3b). The significant diminution of adhesions on dissolved ACES leads suggests successful chronic inhabitance in the polymer mesh and effective ACES cavitation prior to removal (Supplemental Figure 3c,d,e). Histological analysis of cross and longitudinal sections of the nerve demonstrated no significant foreign body response (Supplemental Figure 4) and efficacious repair. In both injectable (c) and cuff (d) embodiments, cavities with clean margins where the electrodes resided can be seen, confirming the mechanism of function of the scaffold. (Supplemental Figure 3). Together, these data suggest that the ACES and its triggerable removal system confer the rehabilitative benefits of ES *and* facilitate a trauma-free removal.

### ACES in minimally-invasive percutaneous implantation

The utility of ACES to stabilize leads and conduct stimulus for deeper anatomic targets was tested by percutaneously implanting electrodes near the femoral and cervical vagus nerves. Under ultrasound guidance, a 14-gauge needle was advanced to the visualized hyperechoic femoral or vagus nerve in anesthetized swine (n = 2). The electrode was placed through an introducer needle to the nerve followed by 1) no injection, 2) 3mL saline injection or 3) 3mL injectable ACES (Figure 4a). The strength of contraction elicited in the distal muscle or nerve was measured in response to stimulation applied at distances of 2 - 28mm from the nerve to assess stabilization and stimulation transmission (Figure 4a-c). Under imaging, the gel and saline hydrodissected the neuromuscular fascial plane. However, saline dissipated quickly, contributing to a decline in stimulus transmission, whereas the gel maintained its shape for the duration of the experiment. Both gel and saline were hypoechoic, though depending on microbubble formation within the gel, varying degrees of echogenicity were visualized, aiding image-guided placement. In the dry condition, nearby off-target tissues were sometimes activated. However, in the presence of the ACES, stimulation evoked significantly higher EMG amplitudes in the target biceps femoris or distal vagus nerve (*p* < 0.05, Student’s two-tailed heteroscedastic t-test). The ACES scaffold enabled PNS even when positioned more than 15 mm from the nerve, while the dry and saline conditions did not evoke a response (Figure 4d,e). Upon repeated actuation, the muscles demonstrated less than 10% reduction in force with over 25 pulses, suggesting no short-term neuromuscular fatigue or reduction in the gel’s conductive capabilities. Upon euthanasia and dissection of the region, a semi-solid and connected path formed by ACES between the lead and the target nerve was observed, substantiating the existence of a conductive pathway enabling the observed stimulation-response results.

**Figure 4.**
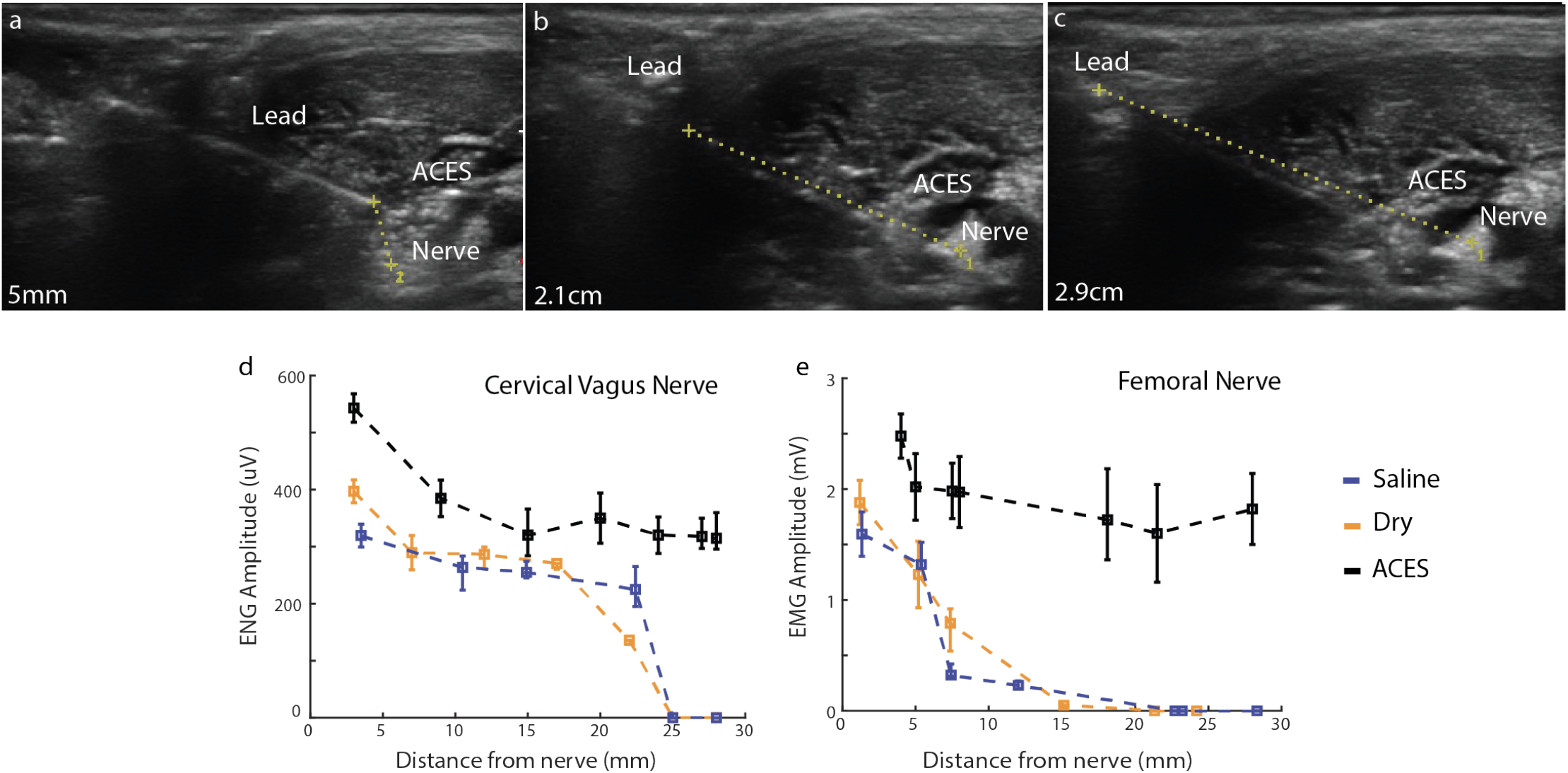
ACES in minimally invasive percutaneous implantation. ACES is injected near the nerve to stabilize the lead and conform to the surrounding geometry as defined by fascial planes. Representative ultrasound imaging from leads positioned at a) 5mm, b) 2.1cm, and c) 2.9cm from the target nerve. Stimulation is significantly higher with ACES bridging the conductive gap (black) as compared to the saline (blue) or dry (orange) conditions for all distances in the d) cervical vagus and e) femoral nerves. ACES can bridge up to a 2.5cm gap.

## Discussion

Although PNS remains a promising therapeutic modality, clinical translation lags behind scientific preclinical advances in part due to the difficulty in lead placement and removal. The ACES platform demonstrates strong potential for hassle-free, quick, and safe placement of leads as well as triggerable release with minimal tissue damage. By minimizing the number of adhesions to the electrode surfaces, the current amplitudes required to activate neural tissues can also be minimized. By facilitating stable electrode placement and preventing lead migration, ACES makes PNS efficacious, enabling improved sensory and motor function following nerve repair.

The form factor and integration of PNS leads in the cuff embodiment of the ACES appreciably builds on the state of art, which are hollow nerve wraps, comprised of collagen or decellularized human nerve allograft that mechanically secure the site of coaptation(*25*). While these wraps preserve the inherent structure of the extracellular matrix (ECM), and reinforce the coaptation site, they provide no targeted acceleration of nerve regeneration or a mechanism to prevent the disuse atrophy and Wallerian degradation that often outpace regeneration. Having leads positioned on both proximal and distal aspects of the nerve repair site aids both directed axonogenesis *and* distal muscle stimulation.

Additionally, ACES enables the neuromodulation of deep-set visceral autonomic nerve targets, which remain difficult to treat by conventional approaches. Historically, deep nerves have rarely been targeted given the degree of risk and morbidity associated with implantation and removal in open surgery. Instead, key nerves such as the celiac and superior mesenteric artery plexus, which critically influence gastric motility and pain, have been treated by either interventional ablation (celiac axis neurolysis) or endovascular ablation. With ACES, these nerves can be accessed via CT and/or ultrasound and the scaffolding will conform to the local anatomical features. Given the lack of histological changes to surrounding muscular bundles and vascular structures, providing electrical stimulations via the gel appears to be safe and generally tolerated. Furthermore, given limitations in imaging resolution, potential gaps in lead placement or lead migration within the scaffold are possible; these issues can be compensated for by the conductive property of the scaffold. This enables minimally invasive image-guided placement, availing a wide range of applications.

Future studies will validate the ACES in larger animal models and other nerve targets. The hydrogel could also be loaded with drugs, growth factors, and immune/neuronal modulators to add a pharmacologic dimension to the intervention. Beyond leads, ACES could be adapted to stabilize other hardware requiring temporary implantation and removal without tissue damage (e.g., catheters, pumps, expanders, depots), enabling new therapeutic interventions previously stifled by hardware implantation and removal challenges.

## Competing Interests

S.S., L.G., P.K, A.S., R.L., G.T. report a provisional patent application (63/240,061) describing the ACES system. Complete details of all relationships for profit and not for profit for G.T. can be found at the following link: https://dropbox.com/sh/szi7vnr4a2ajb56/AABs5N5i0q9AfT1IqIJAE-T5a?dl=0. Complete details for R.L. can be found at the following link: https://dropbox.com/s/yc3xqb5s8s94v7x/Rev&%20Langer&%20COI.pdf?dl=0. A.S is chief medical officer and on the board of CareSignal Health, a digital health company, unrelated to the current work. He is supported by the RSNA Resident Research Award. All other authors declared that they have no competing interests related to the work reported here.

Data and materials availability: All data associated with this study are in the paper or the Supplementary Materials.

## Acknowledgements

Thanks to the Swanson Biotech Core for assistance with histology and Dr. Jaime Cheah for assistance with biocompatibility testing. We thank Declan Gwynne for his help in 3D printing parts for in vivo experiments. This work was supported in part by the Department of Mechanical Engineering and the Karl van Tassel (1925) career development chair, MIT. Original artwork and Illustrations by Virginia E. Fulford, Alar Illustration. We thank the K. Lisa Yang Center for Bionics at MIT for funding support.

## Supplemental Materials

### Supplemental Note 1

*The U*.*S. Food and Drug Administration’s (FDA) Manufacturer and User Facility Device Experience (MAUDE) database indicates that the most common major complications reported for peripheral nerve stimulators, pacemakers, and implanted leads, are lead migration, retained leads, scarring, difficulty of removal, pain, and discomfort (Supplemental Note 1). For example, lead erosion and migration in the StimQ Peripheral Nerve Stimulator System*^19^, *Vertiflex Inc. Superion Interspinous Spacer Prosthesis*^20^, *and Boston Scientific Spinal Cord Stimulators*^21^ *resulted in swelling, discomfort, pain, and/or skin inflammation, requiring removal of hardware without recourse. Scar tissue build-up on electrodes in the StimQ Peripheral Nerve Stimulator System*^19^, *Spinal Modulation Inc. Axium Kit*^22^, *and the Boston Scientific Infinion CX Lead Stimulator*^23^ *led to failures to provide adequate stimulation, pain, and difficulties removing hardware. Pacemaker leads were increasingly left abandoned in place because they could not be removed safely, with one hundred eight-nine pacemaker leads abandoned in situ in 152 patients over a 20-year period*^24,25^

### Supplemental Movie 1

https://www.dropbox.com/s/kuse7nvvqu1fx4h/Nerve&%20graft&%20demo.mp4?dl=0

### Supplemental Movie 2

https://www.dropbox.com/s/4bx5n0akvitb64b/Video&%20Aug&%2005&%2C&%205&%2024&%2036&%20PM.mov?dl=0

**Supplemental Figure 1.**
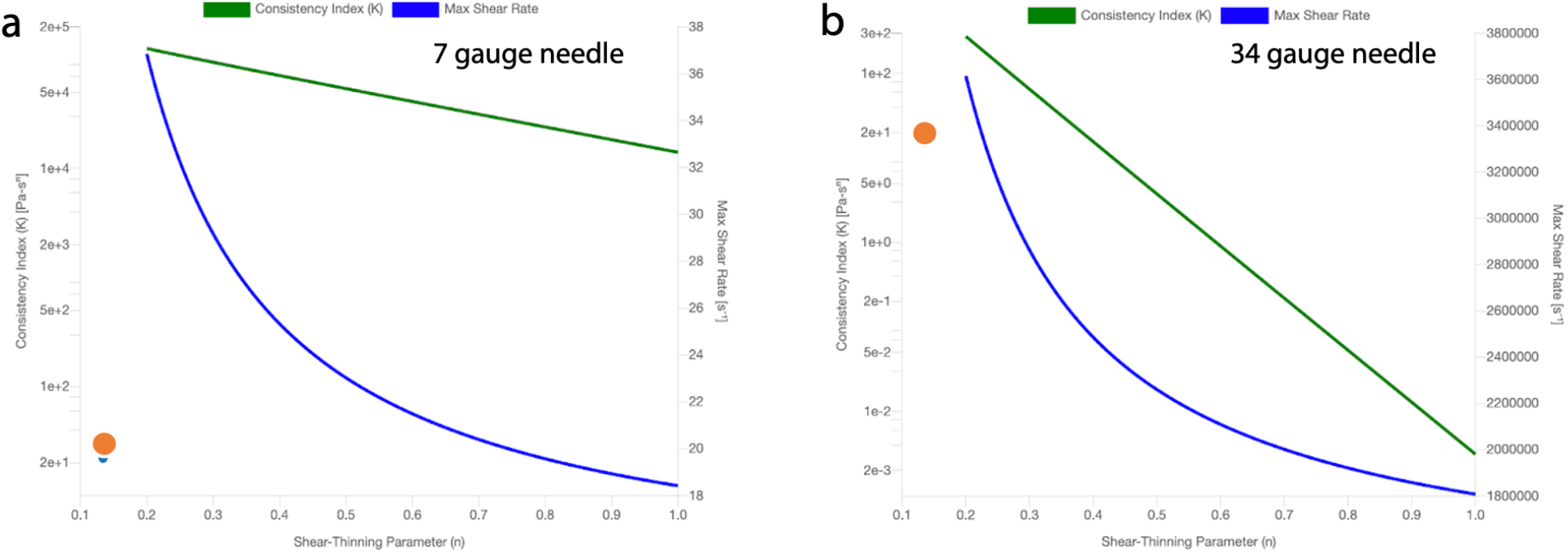
ACES injectable embodiment. Using the gel’s consistency index K = 21.996 Pa*s^n^ and a shear-thinning index (n) of 0.134, injectability was calculated for needle gauges ranging from a) 7 gauge to b) 34 gauge. In all cases, the gel (orange dot) was determined to be injectable.

**Supplemental Figure 2.**
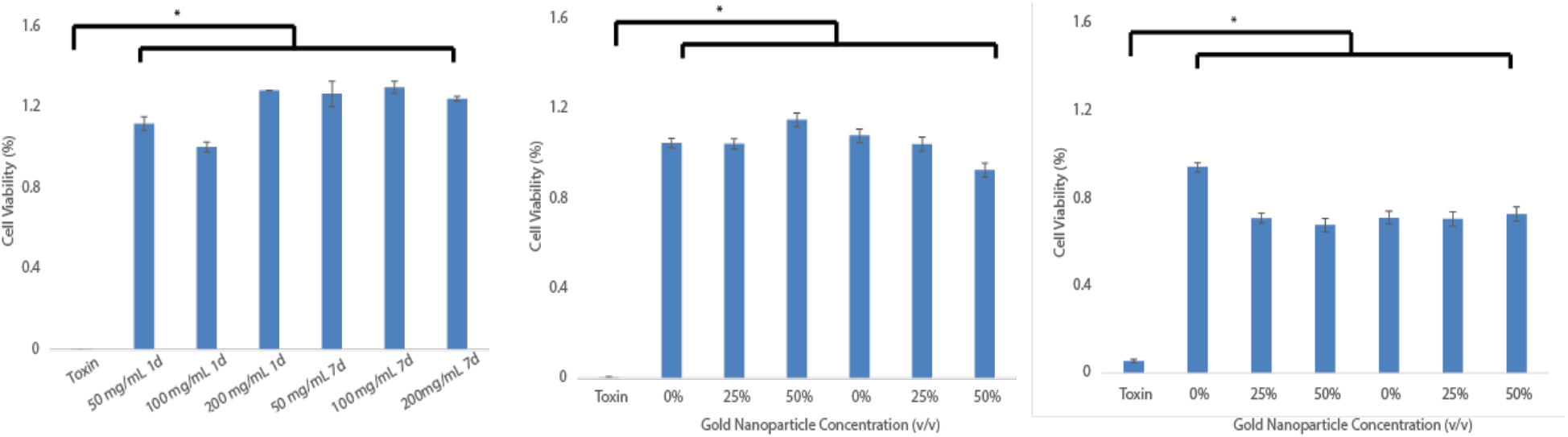
Cell Viability of C2C12 cells demonstrated by an extract exposure test. A) Viability of C2Cl2 cell cultures exposed for a) 1 or 7 days to extracts of given concentrations of alginate and polyacrylamide IPNs b) 24 hours to extracts from formulations of AuNPs at varying concentrations and c) 7 days to extracts from formulations of AuNPs at varying concentrations.

**Supplemental Figure 3.**
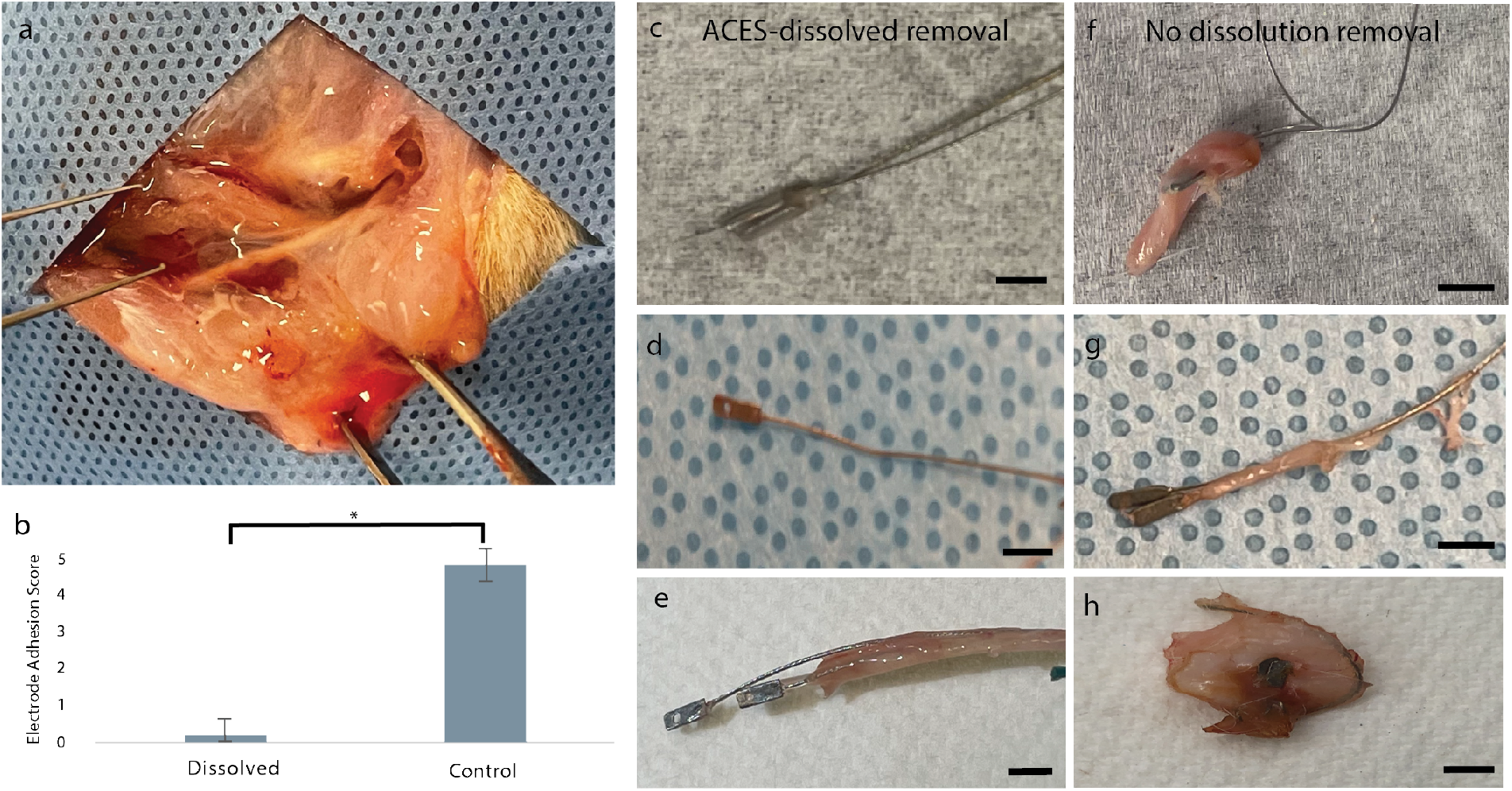
Lead adhesions. a) Representative example of the healed nerve repair after dissolution of ACES. b) Electrode adhesion scores assessed by a blinded reviewer are significantly lower in the dissolved group (p < 0.01, student’s t-test) as compared to the controls. c) Representative images of adhesions on electrodes removed from the experimental (c,d,e) and control (f,g,h) groups in which the ACES was dissolved or not dissolved prior to removal. Adhesions were found to a greater degree in the control group in which the scaffold was not dissolved prior to removal, suggesting greater trauma and disruption of tissues during the removal process. In c, fibrotic tissue can be distinctly seen on the leads, but not on the electrodes which were encompassed by the ACES scaffold. Scale bars represent 10mm.

**Supplemental Figure 4.**
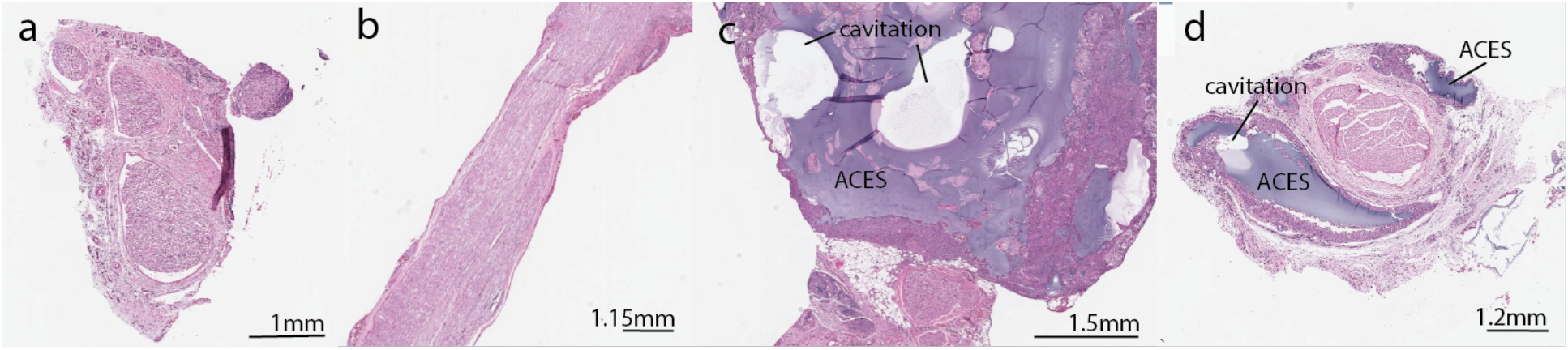
Histological analysis of. a) cross and b) longitudinal sections of the nerve demonstrated no significant foreign body response. C) The ACES gel is visualized in the clear purple in an injectable embodiment. Two cavities with clean margins are visualized where the leads likely resided prior to removal.

**Supplemental Figure 5.**
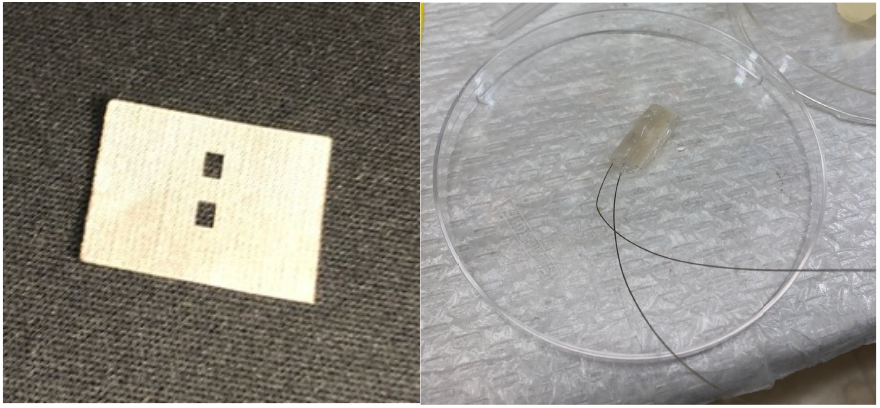
(Left) Vicryl woven mesh was laser cut to form the visualized geometry. (Right) When folded and incorporated into the mold, the windows expose the electrodes.

**Supplemental Figure 6.**
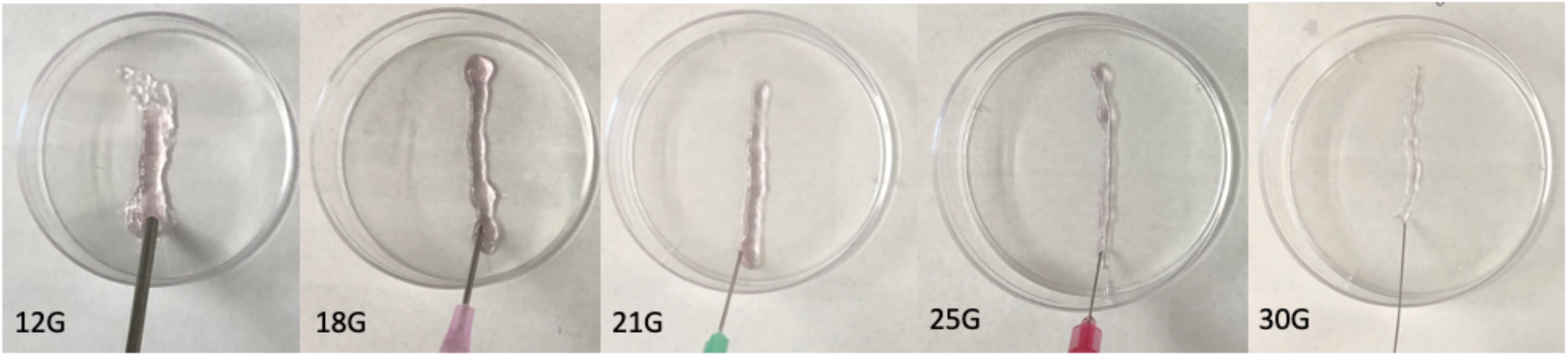
The injectable embodiment of ACES was injected through nozzle gauges ranging from 12 – 30g. In all cases, the gel was extrudable without excessive shear or loss of performance properties.

**Supplemental Figure 7.**
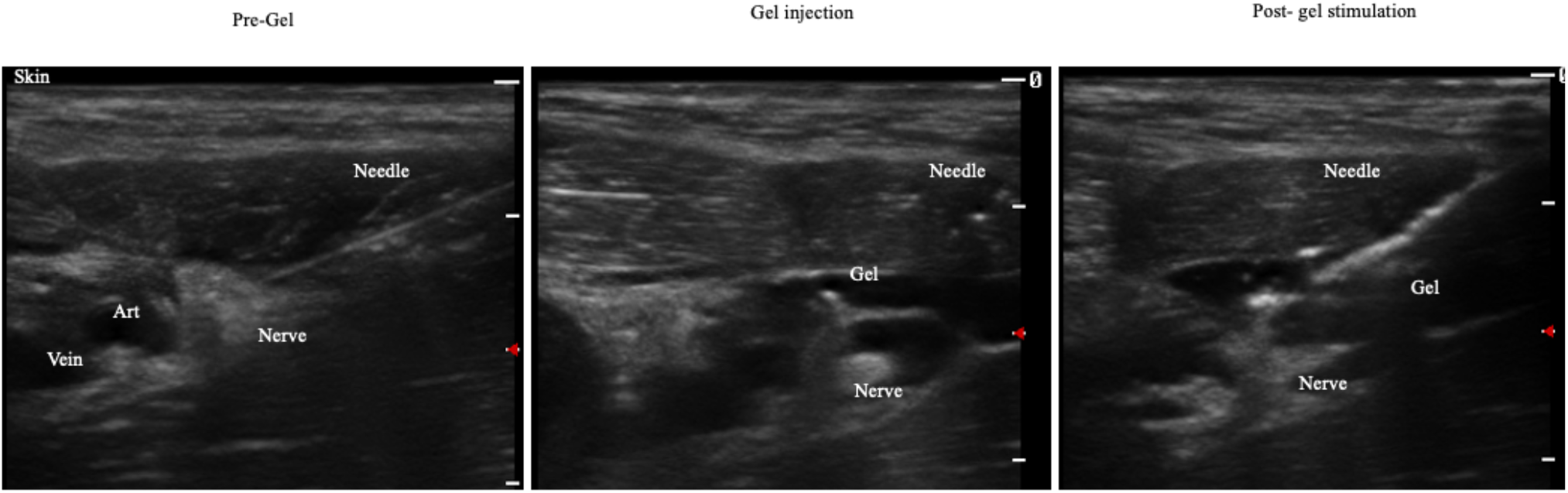
Ultrasound images demonstrates the percutaneous placement of leads with ACES.

**Supplemental Figure 8.**
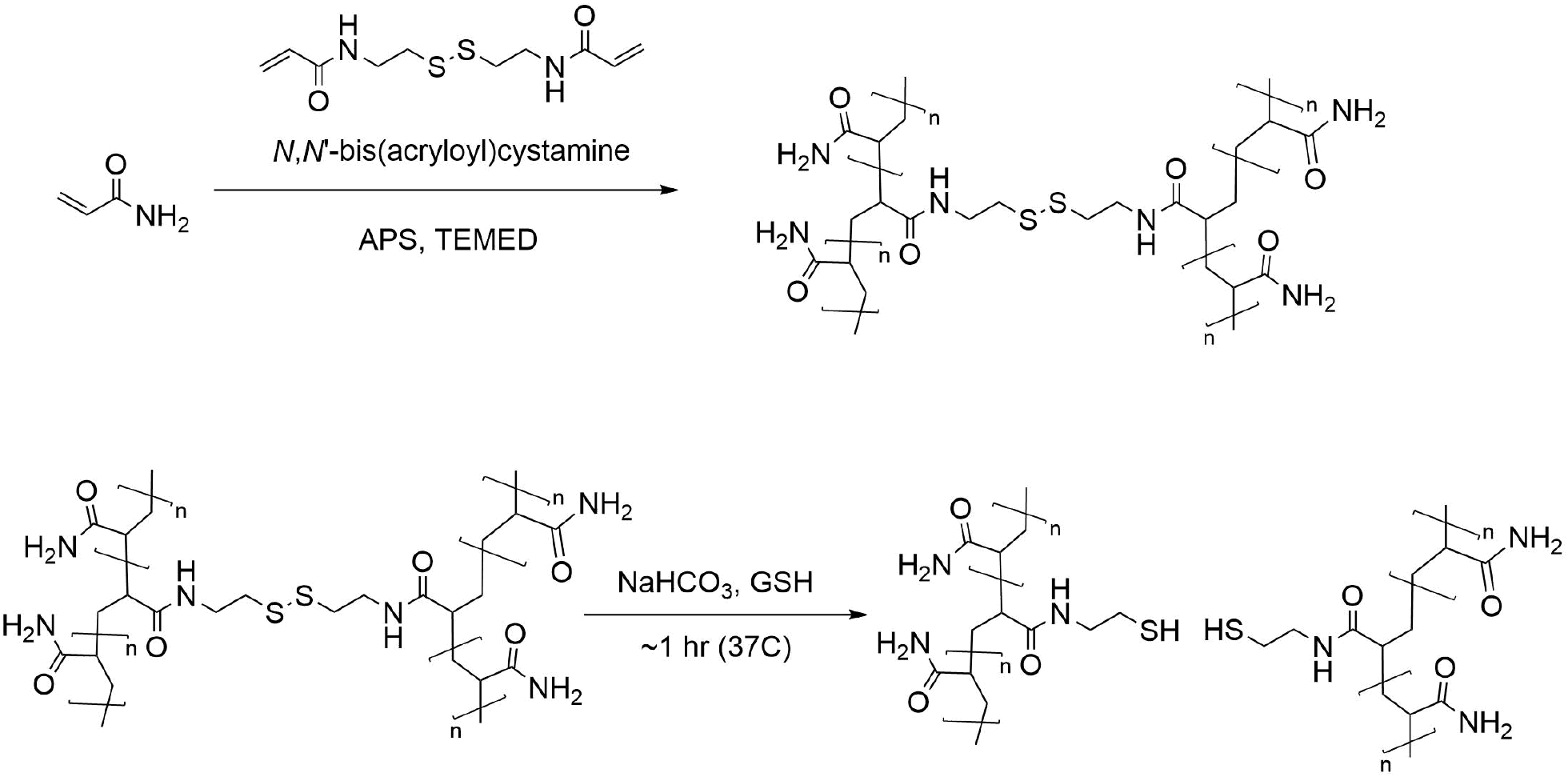
Acrylamide polymerization, crosslinking, and de-crosslinking schema

## Methods

### Formulation of tough hydrogel for adhesion and triggered release of implanted hardware

To form pAm/Alg hydrogels, we first dissolved low viscosity Sodium Alginate (Alg) and Acrylamide (Am) in DI water by vortexing to a final concentration of 8.91mg/mL Alginate and 32.4mg/mL Acrylamide. Then, using a probe sonicator, we dissolved the covalent crosslinker, N’N-bis(acryloyl) cystamine (BAC) in the prepolymer solution at a concentration of 1.08 mg/mL. Following successful dissolution, the Alg/Am/BAC solution was degassed by bath sonication for 20 minutes. Afterwards, radical polymerization initiators Tetramethylethylenediamine (TEMED) and Ammonium Persulfate (APS) and the prepolymer mixture were chilled separately at -20 Celsius for 5 minutes to slow the rate of covalent crosslinking. The chilled APS was added to the Alg/Am/BAC mixture to reach a concentration of 1.56mg/mL, after which the solution was poured into a PDMS mold. At this point, the chilled TEMED was added to the prepolymer mixture to reach a concentration of 0.1 % (v/v) and gently mixed by pipette pumping. The mold was then covered, and the polymerization reaction was allowed to take place at room temperature for 24 hours. Following this interval, the resulting pAm/Alg gel was washed by dialysis for 24 hours to remove unreacted monomer and radical initiator. Alginate crosslinking was achieved by submerging the gel in a 2M solution of calcium chloride for at least 24 hours, resulting in a pAm/Alg IPN. For injectable embodiments, crosslinking was performed *in situ* after injection of the gel. Dual-syringe systems were intentionally avoided to prevent premature crosslinking.

A MS303-76H subcutaneous SU1 stainless steel electrode with a 0.125 inch pad and 10inch wire (Plastics One) was used incorporated into the mold on top of a custom made vicryl mesh (Supplemental Figure 5), cut to the desired shape using a laser cutter (90% speed and 100% power). We incorporated various concentrations of Au-NPs between 0 – 5.5e13 particles/mL (40nm Au NP, Sigma) and used the four-point probe test (Ossila) to determine the most conductive gel concentration.

### Electrode removal assay

To characterize pre-dissolution strength, the wire-embedded IPNs created in a standardized mold were affixed to an Instron apparatus by clamping the electrode lead via the vice grip attachment. The Instron apparatus was programmed to exert upwards pull at a rate of 100mm/minute, and the maximum force required to physically separate the wire from the IPN was observed. Post-dissolution strength on samples incubated in 2mL of dissolution solution (0.5 mg/mL sodium bicarbonate, 0.5 mg/mL reduced glutathione) for 30 minutes.

### Oscillatory rheological measurements and determination of injectability

Time, angular frequency, and flow sweeps were performed on a Discovery Hybrid Rheometer (HR 10, TA Instruments, USA) using am 8.0mm parallel plate, 1mm gap, at 37 °C. Data was analyzed using TRIOS (TA Instruments, USA) Gels were prepared as described above and deposited directly onto the base of the rheometer for at least three trials for each test. A steady shear test was performed to extract the consistency index (K) and shear-thinning index (n) for the material. These were incorporated into an extrudability calculator to determine the maximal needle gauge that would viable for use with the ACES system^31^. Extrudability results were also empirically validated by observing the gel’s cohesion following extrusion through needles gauged 8 – 34.

### Incubation Assay

Preformed and injectable ACES material samples with embedded electrodes were incubated in saline for 4 weeks at 37°C. Mechanical properties were evaluated prior to and after incubation using the electrode removal assay described above.

### Biocompatibility

We performed an extract exposure^32^ test following ISO norms to evaluate the toxicity of the scaffold material. The polymer was formulated at various concentrations (0, 50, 100, 200mg/mL): 0.2g of a polymer block was added to 1mL of DMEM culture medium (Gibco, USA) and stirred at 37 °C for 24 hours or 7 days to create the extract samples. Then, 100uL of fetal bovine serum was added to each 900uL aliquot of the samples and used to treat C2C12 cells plated in a 96 well plate in triplicate. A negative control of untreated media and positive control of media containing MG-132 at a known concentration were also performed. At 24 and 72 hours, cell viability was measured through quantitation of ATP using CellTiter-Glo (Promega, USA) using a Tecan M1000Pro (Tecan Group, CHE).

### In Vivo testing

All animal surgeries were reviewed and approved by the Committee on Animal Care at the Massachusetts Institute of Technology. Male and female Fischer rats (approximately 250g; Charles River Laboratories) were used for all *in vivo* studies. To evaluate the ACES in an open surgical method, the rat model of sciatic nerve transection and repair was selected, given its standardization in the field and known characterization methods^7,8^ and carried out in n = 24 rats.

Animals were premedicated using slow-release buprenorphine and meloxicam. During procedures, animals were anesthetized using 1.5-3% isoflurane in oxygen.

### Open Surgical Approach: Sciatic Nerve Transection and Repair Model

A linear incision was made in the lateral aspect of the hind limb distal to the hip joint. Blunt dissection through the muscle plane between the biceps femoris revealed the sciatic nerve proximal to its branching into the tibial, peroneal, and sural nerves. The nerve was transected using a flat 5 blade and repaired using two 8-0 epineural sutures just proximal to the trifurcation. In n = 18 animals, the embodiment of a nerve cuff was used—the scaffold was inserted and opened underneath the nerve and secured circumferentially around the nerve using 6-0 absorbable sutures. One electrode was carefully positioned distal and proximal to the site of repair. In n = 6 animals, the injectable gel was used wherein the leads were placed at in the proximal and distal positions, as desired. Then, 0.3 – 0.5 mL of gel was injected using a 20 gauge needle on the site. Then, less than 1mL of 2M CaCl was added to the gel, drop by drop. After 5 seconds, excess fluid was absorbed with an absorbent pad or gauze. A microcatheter was placed alongside the electrodes in a position that would allow for local injection of the dissolution solution to the region hosting the electrodes.

Leads were looped to provide a stress/strain loop and tunnel through a subcutaneous conduit in the back. A separate 1 cm incision was made at the nape to host the leads/connector. A custom 3d printed skin port was sutured transcutaneously using 4-0 nonabsorbable suture. This secured the connector to the leads. A silicone cap (McMaster Carr) was used to close the port when not in use. The process is illustrated in Figure 3A – E.

### Rehabilitation

7 animals in whom cuffs were placed were randomly assigned to the electrical stimulation group. Electrical stimulation was administered for rehabilitation every other day for 6 weeks using the Intan RHS Recording/Stimulation System (Intan Technologies, USA). Current was delivered in 30-minute sessions with a 20 hz frequency, 2 mA amplitude, and 100 µs pulse width. Pulses were delivered through the device electrodes in an alternating pattern, with a 100 ms gap between the pulses delivered to the proximal and distal electrodes. All rats were anesthetized, and a ground electrode was placed in the tibialis anterior muscle.

### Electrophysiology

For n = 14 animals with cuff scaffolds, baseline EMG response to electrical stimulation of the TA and GL muscles was recorded and processed 1 week preoperatively, and each week under anesthesia postoperatively for 6 weeks using the Intan RHS hardware and RHX software (Intan Technologies, USA). Current was delivered through the proximal device electrode at 1 Hz and 40 Hz with the following parameters. The distal electrode was grounded, and a ground electrode was also placed subcutaneously over the biceps femoris. EMG data was recorded via bipolar 30G needle electrodes placed in the tibialis anterior and gastrocnemius muscles. Signals were processed using a low pass filter and rectified, and then quantified to identify the 1) threshold at which stimulation elicited a response and 2) the amplitude of the elicited compound muscle activation potentials.

**Supplemental Table 1.** Electrical Stimulation Parameters

| Parameter | 1 Hz Test | 40 Hz test |
| --- | --- | --- |
| Amplitude Ramp | 0 - 7.5 mA at 0.5 mA steps | 0 - 2.5 mA at 0.5 mA steps |
| Frequency | 1 Hz | 40 Hz |
| Pulse Width | 100 us | 100 us |
| Pulse Train Length | 5 s | 300 ms |
| Time Between Pulses | - | 1 s |
| Number of Pulse Trains | 1 | 3 |
| Rest between current steps | 10s | 10s |

### Sensitivity Test

Following a nerve repair, the reinnervation of cutaneous and nociceptive fibers is a key outcome and current limitation of nerve repair. To evaluate the sensory reinnervation, a sensitivity assay was performed: Calibrated forceps were utilized to identify the nociceptive threshold as per previously established methods^33^. Briefly, animals were brought to the experimental room at least 30 min before experimentation and testing was performed during the light phase. The rat was be placed on the bench and loosely restrained using a towel to cover the eyes and prevent environmental stimulation. The tips of the forceps were be positioned on the paw. Force was manually incremented at a rate of 20 g s−1 until the limb was withdrawn. The threshold was be noted and the rat was allowed to rest for approximately a minute. Measurements were repeated five times and the averages were compared between groups using two-tailed heteroscedastic t-tests.

### Electrode Removal Force Assay

Dissolution was performed in half of the animals with cuff and injectable scaffolds and those undergoing electrical stimulation (n= 13), while the rest were assigned to a control non-dissolution group (n=13). For dissolution, a solution of 0.5M GSH, 0.5M SBC, and 1M EDTA were injected into the space underneath biceps femoris and incubated for 30 minutes to promote dissolution of ACES. A small subcutaneous incision was performed to locate the electrode and attach it to the suction grips of an Instron tensile testing machine or handheld tensiometer. A custom positioning pin aligned the direction of the tension parallel to the sagittal plane of the animal to enable a pulling force that would allow the electrode leads to slide through existing muscle plane boundaries.

### Tissue morphology assessment

A blinded plastic surgeon scored adhesions and fibrotic response to the hardware on the nerve, adhesions to electrodes after removal, and fibrotic response to the scaffolds on a scale of 0 – 5, where a greater number represented increased scarring.

### Tissue Harvest

Following euthanasia, the sciatic nerve, its branches, tibialis anterior, and gastrocnemius were carefully harvested from both sides of each animal. The mass of each muscle was recorded, and ratios were calculated between affected and contralateral unaffected limbs in each animal for each muscle. All tissues were fixed in 4% paraformaldehyde, paraffin processed, embedded, and then sectioned. Tissues were stained with 1) hematotoxylin and eosin to study general structure, regeneration, and trauma and 2) trichrome to assess fibrotic responses. A semi-automated script in ImageJ was used to identify the total number of axons 0.5cm proximal to the site where the scaffold was implanted. Axon count 1mm distal to the branching point and at 1.5 cm distal to the branching point.

### Percutaneous Placement of ACES in Swine

To evaluate the capabilities of ACES in facilitating leads in a minimally-invasive manner, a model of ultrasound-guided placement at the femoral nerves was performed. A 14-gauge needle was advanced to the target nerve, visualized using a linear ultrasound probe (SonoScape S9 portable ultrasound system, Model ST-180, SonosScape Medical Corp, CHN) at a depth of 4 inches. 3mL of the ACES material or saline (control) was injected. Then, an electrode was advanced through the needle and placed at a measured distances from the nerve. Electrophysiology of the biceps femoris or distal vagus nerve were performed using 2 bipolar 32-gauge needle electrodes (Natus Medical, USA). Stimulation was performed using a model 2100 isolated pulse stimulator (A-M Systems, USA) at 2 Hz, 20 pulses, 2-6mA.

